# Kinesin family member Kif23 regulates cytokinetic division and maintains neural stem/progenitor cell pool in the developing neocortex

**DOI:** 10.1101/2023.10.27.564302

**Authors:** Sharmin Naher, Takako Kikkawa, Kenji Iemura, Satoshi Miyashita, Mikio Hoshino, Kozo Tanaka, Shinsuke Niwa, Jin-Wu Tsai, Noriko Osumi

## Abstract

Accurate mitotic division of neural stem cell/progenitor cells (NSPCs) is crucial for the coordinated generation of progenitors and neurons in the developing cortex. Here, we investigated the pivotal role of Kif23, an N-kinesin motor protein, in embryonic mouse NSPCs. We found that Kif23 is highly expressed in the mitotic NSPCs within the embryonic cortex of both mouse and human. Knockdown (KD) of *Kif23* led to precocious neurogenesis, attributed to an accelerated cell cycle exit, likely resulting from disrupted mitotic spindle orientation and impaired cytokinesis. *Kif23* KD induced upregulation of the γ-H2AX-p53-p21 signaling pathway, ultimately culminating in cytokinetic failure. Additionally, Kif23 depletion perturbed the apical surface structure of NSPCs and disrupted the proper localization of apical junctional proteins. Importantly, we demonstrated the successful rescue of *Kif23* KD-induced phenotypes by introducing wild-type human *KIF23*, but not by a variant of *KIF23* with a microcephaly-associated mutation. Our findings underscore the critical role of Kif23 in cortical development and provide novel insights into the intricate molecular mechanisms underlying pathogenesis of microcephaly.

**Graphical abstract:** 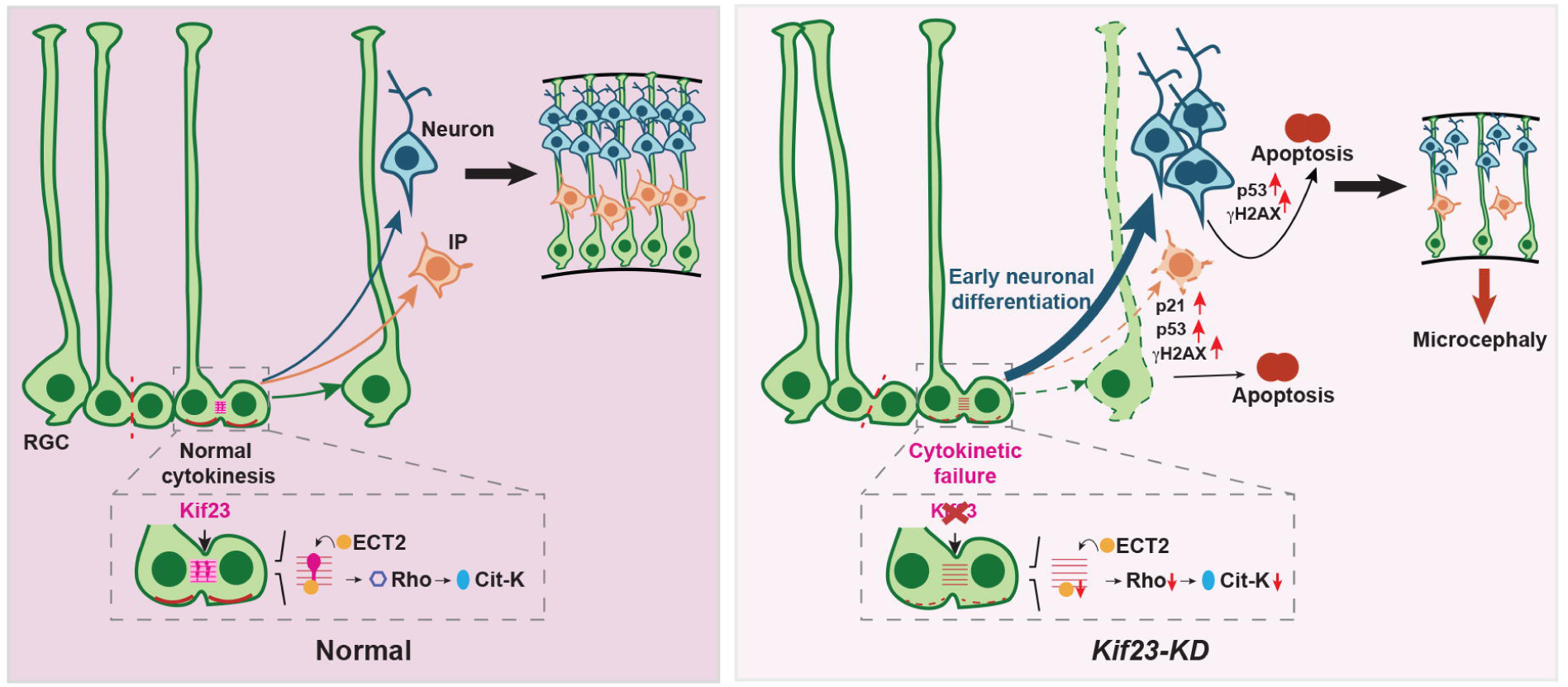

## Introduction

The development of the mammalian cerebral cortex is a highly intricate and meticulously orchestrated series of events. Central to this process are the pivotal roles played by neural stem/progenitor cells (NSPCs), encompassing their initial proliferation and the subsequent neuronal differentiation and migration (Götz & Huttner, 2005). In the early stages of corticogenesis, NSPCs residing in the ventricular zone (VZ) of the cortex engage in numerous rounds of proliferative division, thereby expanding the progenitor pool. As corticogenesis advances, neurogenic division takes precedence, giving rise to neurons either directly or indirectly through transit-amplifying progenitors, including intermediate progenitors (IPs) (Götz & Huttner, 2005; Kriegstein et al., 2006). Disturbance in these finely tuned processes can result in abnormal brain size and structural anomalies.

The Kinesin superfamily motor proteins (Kifs) are responsible for controlling microtubule dynamics and facilitating intracellular transportation of organelles, mRNAs, and protein complexes along microtubules (Hirokawa et al., 2009). In the nervous system, members of Kif family are localized in both mitotic and post-mitotic cells. In post-mitotic neurons, Kifs are essential motor proteins involved in neuronal migration, transport, and synaptic transmission (Joseph et al., 2021). Dysregulated expression of KIFs has been implicated in several neurological disorders, including amyotrophic lateral sclerosis, epilepsy, and Alzheimer’s disease (Asselin et al., 2020; Guillaud et al., 2020; Lucero et al., 2022). While the role of Kifs in neurons is well-established, their functions in NSPCs during embryonic brain development remain less explored.

Within the Kif family, the kinesin-6 subfamily, comprised of three members - Kif20a, Kif20b, and Kif23, is notably recognized for its pivotal role in mitotic processes (Hirokawa et al., 2009). Specifically, Kif20a and Kif20b have emerged as key regulators in orchestrating the division of NSCPs during corticogenesis (Janisch et al., 2013; Geng et al., 2018). Meanwhile, Kif23, a prominent member of the Kif 6 family, assumes a critical role as a major component of the central spindle complex. During mitosis in cell lines, Kif23 conspicuously localizes at the cleavage furrow and the midbody, where it governs the bundling of anti-parallel microtubules and facilitates cytokinesis (Adams et al., 1998; Pavicic-Kaltenbrunner et al., 2007). Furthermore, in cultured neuronal cells, Kif23 emerges as an essential factor in maintaining dendritic morphology and composition (Sharp et al., 1997; Yu et al., 2000). Notably, recent genetic studies in humans have identified mutations in human *KIF23* gene associated with microcephaly (Karaca et al., 2015; Boonsawat et al., 2019). However, the precise role of Kif23 in the intricate development of the mammalian cortex remains uncharacterized.

In this study, we discovered that Kif23 exhibits exclusive expression within the NSPCs residing in the developing neocortex. To elucidate the *in vivo* role of Kif23 in NSPCs, we harnessed the technique of *in-utero* electroporation to targeted knock-down of *Kif23*. This depletion of Kif23 had far-reaching consequences, disturbing the delicate balance of spindle orientation, cytokinesis, and structural integrity of the apical surface. These disruptions resulted in the early depletion of NSPCs, followed by a cascade of events, including neuronal apoptosis and substantial reduction in neuronal production. Notably, our study not only sheds light on the essential functions of Kif23 in NSPCs but also provides valuable insights into a novel mechanism underlying microcephaly. Specifically, we offer a fresh perspective on how the loss of human KIF23 may give rise to microcephaly by illuminating hitherto unexplored mechanism that controls cytokinesis in diving NSPCs throughout cortical development.

## Results

### Kif23 is highly expressed in the embryonic mouse neocortex

In order to elucidate the role of Kif23 in neocortical development, our initial investigation focused on the expression of *Kif23* within the developing neocortex. We leveraged the spatial gene expression dataset derived from the E15.5 mouse brain (Tsai et al., under revision), which categorized gene expression profiles into 12 distinct clusters (Figure 1A, left). Interestingly, our analysis revealed that Kif23 exhibited specific expression within cluster 10, a cluster primarily situated in the ventricular zone (VZ), where NSPCs are enriched (Figure 1A, middle and right). We indeed confirmed the abundance of *Kif23* mRNA in the VZ by *in situ* hybridization at E14.5 (Figure 1B).

**Figure 1.**
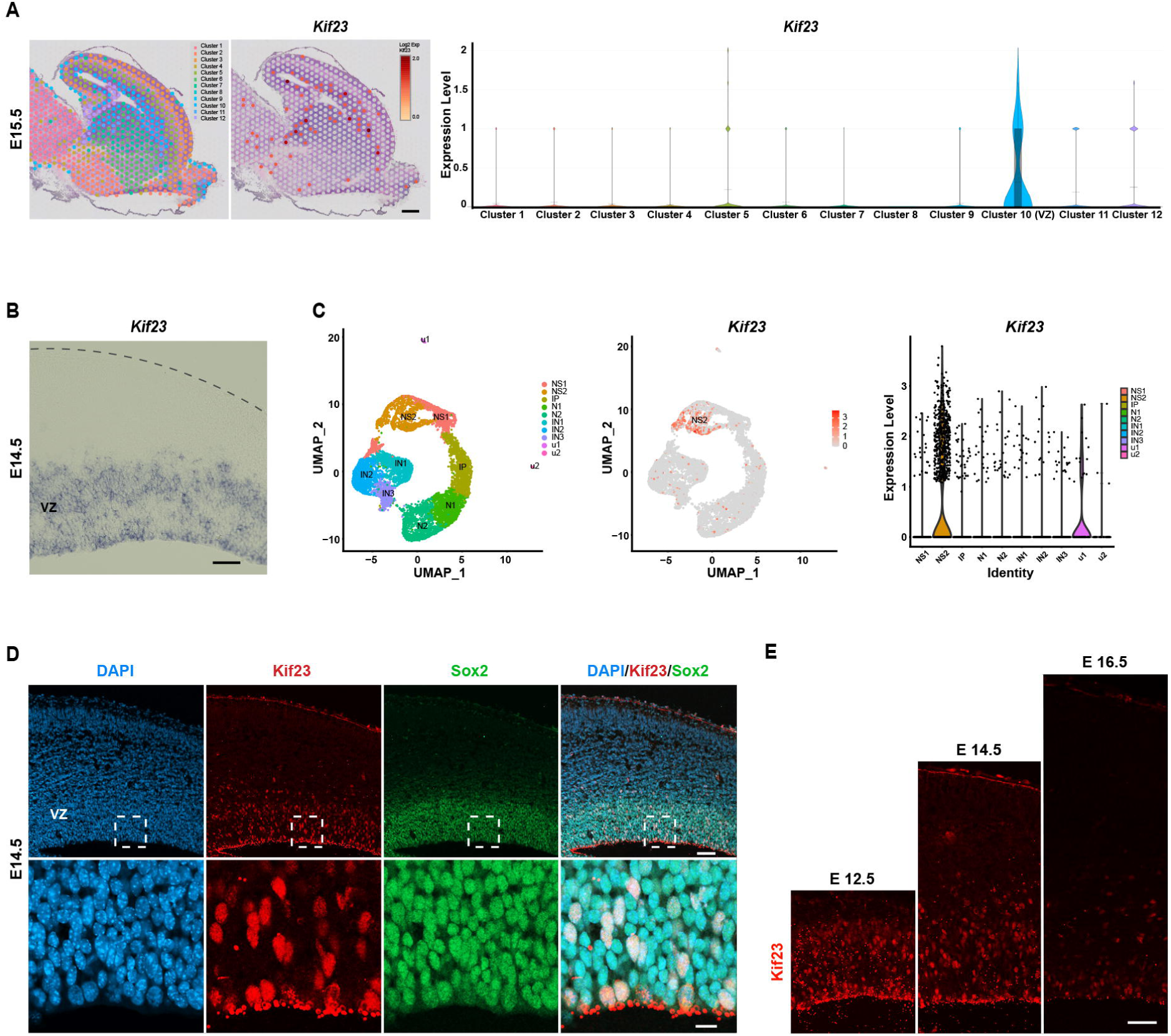
Kif23 expression in the developing neocortex. **(A)** Visium spatial gene expression analysis of E15.5 mouse brain showing gene expression clusters (left), spatial plots (middle), and violin plots (right) to show the spatial expression of *Kif23. Kif23* is enriched in the cluster 10. Scale bar, 300 μm. **(B)** Expression of *Kif23* mRNA detected by *in situ* hybridization of E14.5 embryonic mouse cortex. *Kif23* is localized in the ventricular zone (VZ). Scale bar, 30 μm. **(C)** Single-cell RNA-seq analysis of E14.5 mouse cortex showing cell type clusters (left), feature plots (middle), and violin plots (right) to show the expression of *Kif23. Kif23* is enriched in the neural stem/progenitor cells cluster 2 (NS2). NS1, neural stem/progenitor cells cluster 1; IP, intermediate progenitor cells cluster; N1 and N2, neurons cluster 1 and 2; IN1, IN2 and IN3, interneurons cluster 1, 2 and 3. **(D)** Representative image of E14.5 mouse cortex stained with the antibodies to Kif23 and Sox2, a marker of neural stem progenitor cells (NSPCs) and DAPI. Kif23 is co-localized with Sox2. Boxed areas are magnified in the bottom panel. Scale bars, 50 μm (top) and 10 μm (bottom). **(E)** Immunostaining of the mouse cortex at E12.5, E14.5 and E16.5 showing expression of Kif23 at a higher level in E12.5 and E14.5. Scale bar, 50 μm.

We next examined the specific cell types expressing Kif23 in the embryonic mouse brain using previously published single-cell RNA sequencing (scRNA-seq) data from E14.5 mouse telencephalon (Loo et al., 2018). Through a meticulous reanalysis of this scRNA-seq dataset, we stratified cell clusters into distinct categories: NSPCs (NS), IPs (IP), neurons (N), and interneurons (IN) (Figure 1C, left). Remarkably, our analysis unveiled a notable enrichment of *Kif23* within the actively proliferating NSPCs, particularly prominent in the NS2 cluster expressing *Top2a*, a marker of proliferative cells (Figure 1C, middle and right).

We also confirmed that Kif23 protein expression in the NSPCs by co-immunostaining with Sox2, a marker for NSPCs (Figure 1D). During cortical development, Kif23 protein was detected at a higher level in the neocortical VZ progenitor cells at E12.5 and E14.5 during neurogenic stages, while its expression declined at E16.5 when the neurogenesis decreased (Figure 1E). This stage-specific expression in NSPCs suggests a potential role of Kif23 in regulating early cortical neurogenesis.

### Knockdown of *Kif23* induces premature neuronal differentiation

Next, we employed an *in-utero* electroporation technique to knockdown (KD) *Kif23* by delivering a small interfering RNA (siRNA) vector targeting the mouse *Kif23* gene, together with an EGFP expression vector, into the mouse neocortex at E14.5 (Figure 2A). Successful *Kif23* siRNA-mediated reduction of endogenous Kif23 protein expression was confirmed at E15.5 (Figure 2B). In this context, we examined the distribution of GFP^+^ cells across various zones of the developing cortex two days after *Kif23-*KD (at E16.5). Notably, the percentage of GFP^+^ cells in the VZ/subventricular zone (SVZ) exhibited a significant decrease in *Kif23*-KD cortices compared to the control group, whereas the proportion of GFP^+^ cells in the IZ significantly increased in the *Kif23*-KD cortices (Figures 2C and 2D).

**Figure 2.**
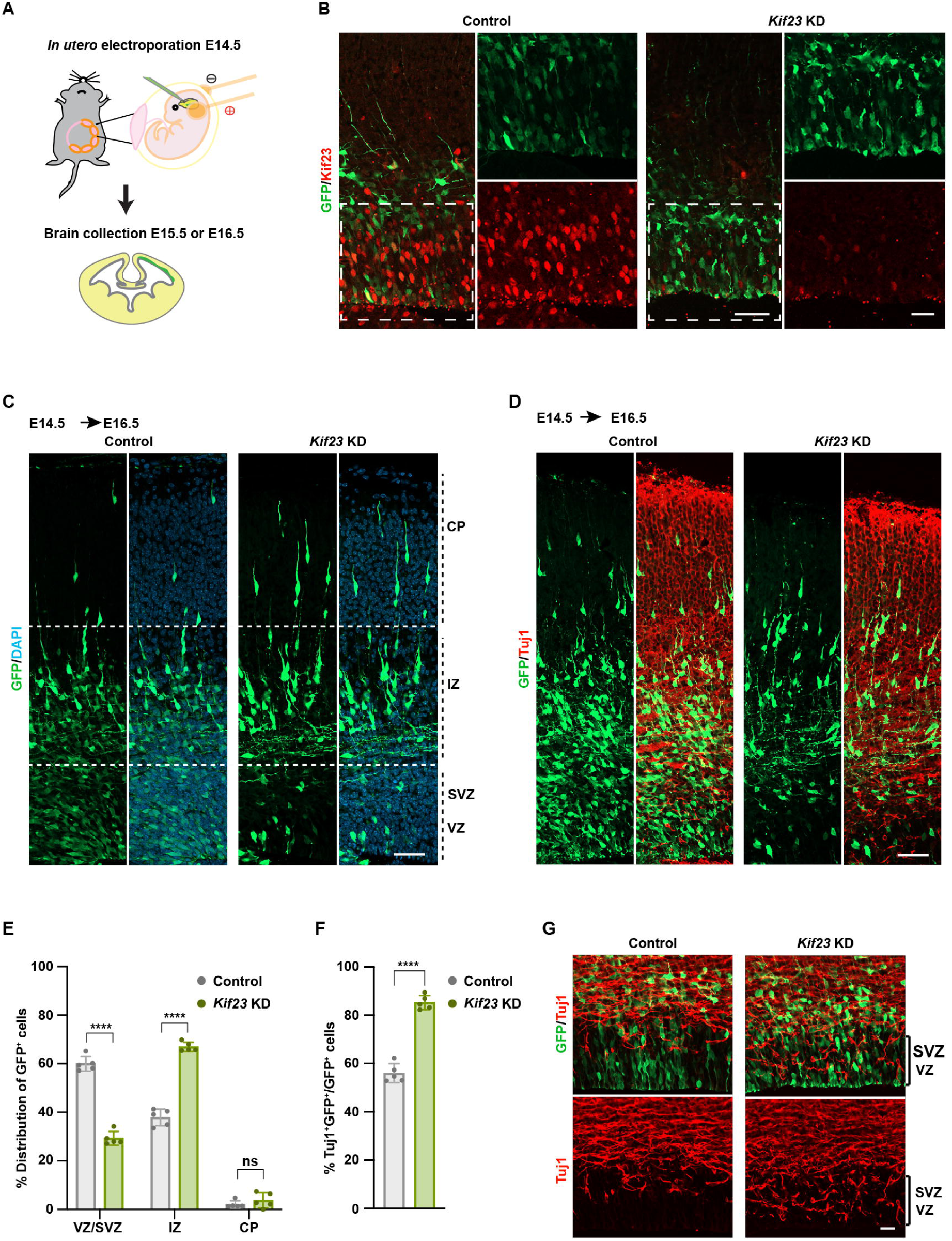
Kif23 knockdown induces premature neuronal differentiation. **(A)** Schematic diagram of *Kif23* knockdown (KD) by *in-utero* electroporation. The embryonic mouse brains were electroporated at E14.5 with control or *Kif23* siRNA. **(B)** Representative images stained for GFP, Kif23, and DAPI in the E15.5 mouse cortices after *in-utero* electroporation. The expression of Kif23 is effectively suppressed by *Kif23* siRNA. Boxed areas are magnified. Scale bars, 50 μm (left) and 25 μm (right). **(C)** Representative images of the distribution of GFP^+^ cells in the control and *Kif23-*KD cortices. Dashed lines illustrate the borders among the ventricular zone/ subventricular zone (VZ/SVZ), intermediate zone (IZ), and cortical plate (CP). Scale bar, 50 μm. **(D)** Representative images of the control and *Kif23-*KD cortices at E16.5 stained for GFP and Tuj1. Scale bar, 50 μm. **(E)** Quantification of the distribution of GFP^+^ cells in VZ/SVZ, IZ, and CP, respectively. The data represent the mean ± SD (n = 5). Two-way ANOVA with Bonferroni’s multiple comparisons test, \*\*\*\**P* < 0.0001; ns, not significant. **(F)** Quantification of the percentage of Tuj1^+^GFP^+^ cells relative to the total GFP^+^ cells. The data represent the mean ± SD (n = 5). Two-tailed Student’s *t*-test, \*\*\*\**P* < 0.0001. **(G)** Representative images of the control and *Kif23-*KD cortices at E15.5 stained for GFP and Tuj1. Scale bar, 20 μm.

In light of the migration of NSPCs in the VZ/SVZ to the IZ and CP upon exiting the cell cycle, the observed perturbation in cell distribution hints at the potential induction of premature differentiation attributable to *Kif23*-KD; we conducted an analysis of the neuronal marker Tuj1 (βⅢ-Tubulin) expression within both the control and *Kif23*-KD groups. Remarkably, a significantly larger portion of GFP^+^ cells in the *Kif23*-KD cortices co-expressed Tuj1 in comparison to the control cortices (Figures 2E and 2F). Furthermore, this trend persisted with the observation of elevated Tuj1 expression within the VZ of *Kif23*-KD cortices at E15.5 (Figure 2G). Collectively, these findings substantiate the notion that *Kif23*-KD fosters premature neuronal differentiation in the developing cortex.

### Knockdown of *Kif23* inhibits proliferation of progenitor cells

Earlier investigations have demonstrated the pivotal role of Kif23 in cell proliferation, especially in the context of cancer cell lines (Liu et al., 2020; Li et al., 2022). Consequently, we sought to elucidate whether the observed increase in differentiated cells could be attributed to changes in the maintenance and proliferation of NSPCs. Notably, our findings revealed that *Kif23-*KD led to a significant decrease in the percentage of cells that co-expressed GFP and Pax6 (a NSPC marker), as compared to the control group (Figures 3A and 3B). Similarly, the proportion of cells positive for both GFP and Tbr2 (an IP marker) was significantly lower in the *Kif23*-KD group in comparison to the control group (Figures 3C and 3D). These observations unequivocally signify a depletion in the progenitor pool induced by *Kif23*-KD.

**Figure 3.**
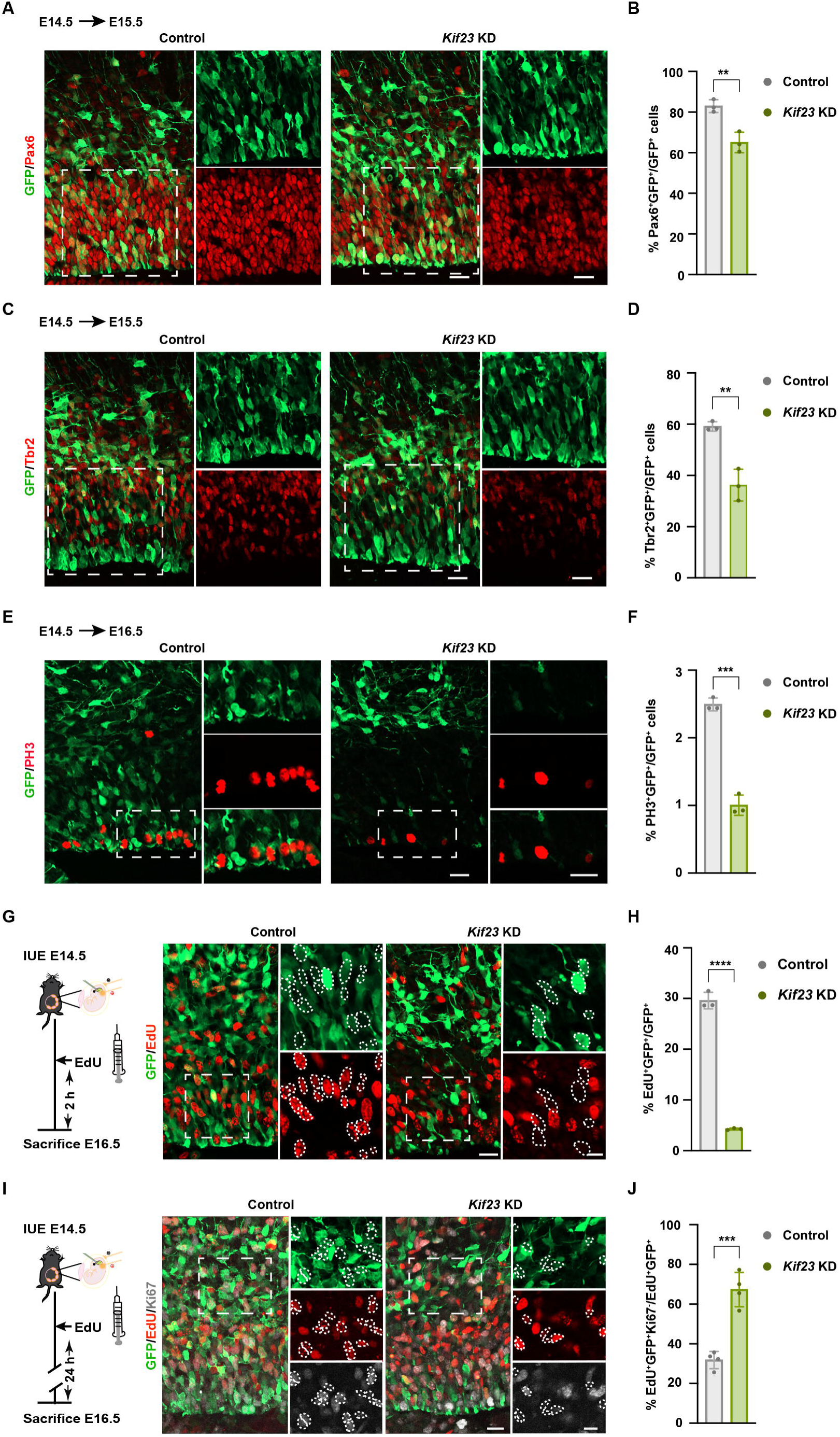
Kif23 deficiency depletes NSPCs pool and induce premature cell cycle exit. **(A, C, and E)** Representative images of the control and *Kif23-*KD cortices stained for GFP and Pax6 **(A),** or GFP and Tbr2 **(C),** or GFP and PH3 **(E)**. Boxed areas denote zoomed areas. Scale bars, 20 μm **(A and C)**; 25 μm **(E)**. **(B, D, and F)** Quantification of the percentage of Pax6^+^GFP^+^ cells **(B)** or Tbr2^+^GFP^+^ cells **(D)** or PH3^+^GFP^+^ cells **(F)** relative to the total GFP^+^ cells. The data represent the mean ± SD (n = 3). Two-tailed Student’s *t*-test, \*\**P* < 0.01; \*\*\**P* < 0.001. **(G and I)** Timeline of the *in-utero* electroporation and EdU injection (left). Electroporated cortical sections were stained for GFP and EdU **(G)** or GFP, Ki67, and EdU **(I)** (right). Boxed areas denote zoomed areas. Scattered lines encircle GFP^+^ cells that are EdU^+^ or Edu^−^ **(G)** or GFP^+^ EdU^+^ cells that are Ki67^+^ or Ki67^−^ **(I)**. Scale bars, 20 μm. **(H and J)** Quantification of the percentage of EdU^+^GFP^+^ cells **(H)** or EdU^+^Ki67^−^GFP^+^ cells **(J)**. The data represent the mean ± SD (n = 3). Two-tailed Student’s *t*-test, \*\*\**P* < 0.001; \*\*\*\**P* < 0.0001.

To gauge the impact of *Kif23*-KD on the proliferation of NSPCs, we conducted an assessment of mitotic activity by staining for the mitotic marker phospho-histone-3 (PH3). Notably, a significant decrease in the number of PH3^+^ cells was observed in *Kif23*-KD cortices when compared to the control group (Figures 3E and 3F). To provide a comprehensive evaluation of the overall proliferative capacity of NSPCs, we adopted a dual approach. Firstly, we electroporated the embryonic mouse cortex at E14.5 and subsequently administrated an intraperitoneal injection of EdU (labeling S-phase proliferating cells) just two hours before the termination of the experiment at E16.5. Remarkably, the percentage of EdU^+^GFP^+^ cells among the GFP^+^ cells, which serves as a reliable proliferative index, exhibited a noteworthy reduction in response to *Kif23*-KD (Figures 3G and 3H). These compelling results collectively underscore the indispensable role of Kif23 in promoting the proliferation of NSPCs during the critical period of embryonic neurogenesis.

It is established that when the proliferation of NSPCs is inhibited, these cells tend to exit the cell cycle and differentiate into neurons (Qiao et al., 2018). We thus examined whether these Kif23-deficient progenitors prematurely departed from the cell cycle. To facilitate this analysis, we performed *in-utero* electroporation of the embryonic mouse cortex at E14.5, followed by an EdU injection at E15.5. Subsequently, we conducted an examination of GFP^+^EdU^+^Ki67^−^ cells, at E16.5, which represent the cell proportion exiting the cell cycle. Our observation uncovered a significant increase in the percentage of GFP^+^EdU^+^Ki67^−^ cells among the GFP^+^EdU^+^ cell population in response to *Kif23*-KD (Figures 3I and 3J). This finding provides clear evidence that the loss of Kif23 served to promote an early exit from the cell cycle. Therefore, our results consistently affirm the critical role of Kif23 in the maintenance and renewal of NSPCs.

### Kif23 regulates mitotic spindle organization and orientation

Given the essential role of Kif23 in central spindle assembly and cytokinesis within HeLa cells (Mishima et al., 2004), we examined the subcellular localization of Kif23 protein across various phases of the cell cycle. The cell cycle phases were determined by assessing nuclear morphology via DAPI staining and the positions of a centrosomal protein γ-tubulin. Our observation revealed distinct patterns of Kif23 localization: during G2, S, and prophase, Kif23 predominantly resided within the nucleus of NSPCs (Figure 4A). However, as the cell cycle progressed, Kif23 transitioned to different cellular compartments: it relocated to the spindle during prometaphase and metaphase, occupied the midzone during anaphase and telophase, and ultimately gathered within the midbody during the cytokinetic phase (Figure 4A).

**Figure 4.**
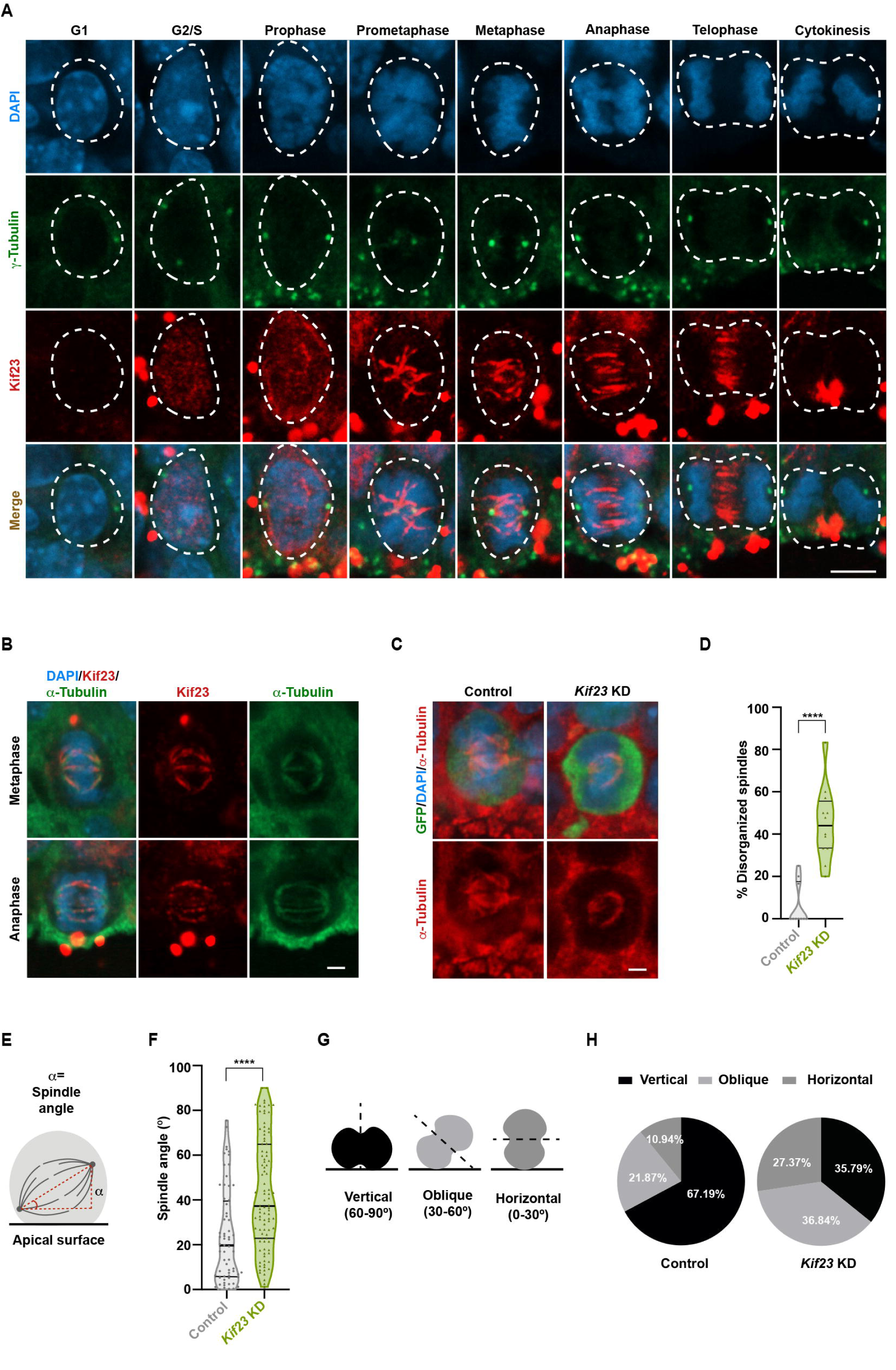
Loss of Kif23 disrupts the organization and orientation of mitotic spindle. **(A and B).** Representative images of WT E14.5 cortex stained for Kif23, DAPI and γ tubulin (centrosomal marker) **(A)** or α-tubulin (microtubule marker) **(B)**. Dashed lines indicate cells in the different stages of the cell cycle. Scale bars, 5 μm **(A)**; 2 μm **(B)**. **(C and D).** Representative images of the electroporated brain sections at E15.5 stained for GFP and α-tubulin **(C)**. Scale bar, 2 μm. Quantification of the percentage of disorganized spindles **(D)**. The data represent the mean ± SD (n = 10 for the control group; n=12 for the *Kif23-*KD). Two-tailed Student’s *t*-test, \*\*\*\**P* < 0.0001. **(E and F).** Graphical representation of spindle angle measurement **(E)**. Distribution of spindle angles (°) **(F)**. The data represent the mean ± SD (n = 10 samples for the control group; n = 12 samples for the *Kif23-*KD group). Two-tailed Student’s *t*-test, \*\*\*\**P* < 0.0001. **(G and H).** Graphical representation of vertical, oblique, and horizontal divisions **(G)**. Black dash lines indicate the cleavage plane. Quantification of the percentage of each class of spindle cleavage plane orientation **(H)**. Data represent n = 64 cells from 10 samples for the control group; n = 95 cells from 12 samples for the *Kif23-*KD group.

The dynamic shifts in Kif23 localization within the NSPCs, particularly its co-localization with spindle microtubule marker α-tubulin during mitosis (Figure 4B), strongly point towards Kif23’s pivotal role in orchestrating mitotic spindle organization. This notion gains support from a significant increase in the prevalence of disorganized spindles among mitotic progenitors following Kif23 depletion (Figures 4C and 4D). Furthermore, our investigation delved into the orientation of the mitotic spindle, scrutinizing the cleavage plane’s alignment in relation to the apical surface (Figure 4E). Notably, *Kif23*-KD NSPCs displayed a larger spindle angle when compared to the control group (Figure 4F). Additionally, the percentage of mitotic progenitors with spindle cleavage planes oriented horizontally (control, 10.94%; *Kif23*-KD, 27.37 %) or obliquely (control, 21.87%; *Kif23*-KD, 36.84 %) in relation to the apical surface increased after *Kif23*-KD, while the percentage of the those with vertically oriented spindle cleavage planes decreased (control, 67.19%; *Kif23*-KD, 35.79 %) (Figures 4G and 4H). These findings collectively underline the impact of Kif23 on the proper orientation of the mitotic spindle, which, when impaired due to Kif23 depletion, may lead to alterations in spindle cleavage plane orientation and, consequently, influence the mode of NSPC division, in line with a previous study (Yingling et al., 2008).

### Kif23 deficiency induces apoptotic cell death

Interestingly, our investigation unveiled a significant increase in number of GFP^+^ cells within the CP and IZ, along with a corresponding decrease in the VZ/SVZ three days after *Kif23*-KD (Figures S1A and S1B). Moreover, a significant reduction in the overall GFP^+^ cell population was evident in the KD group (Figures S1A and S1C). This profound reduction of GFP^+^ cells suggests the involvement of additional underlying mechanisms beyond mere precocious neuronal differentiation. Subsequently, we conducted an examination of apoptosis in the Kif23-deficient cortices one day after electroporation at E15.5. The results were striking, revealing a noteworthy increase in pyknotic nuclei, indicative of chromatin and nucleus condensation during apoptotic cell death, within the VZ, SVZ, and IZ, as detected by DAPI staining (Figures 5A and 5B). Immunostaining with the apoptosis marker, cleaved caspase 3 (CC3), further confirmed our observation, highlighting the heightened apoptotic activity within *Kif23*-KD cortices in comparison to the control group (Figures 5A and 5C).

**Figure 5.**
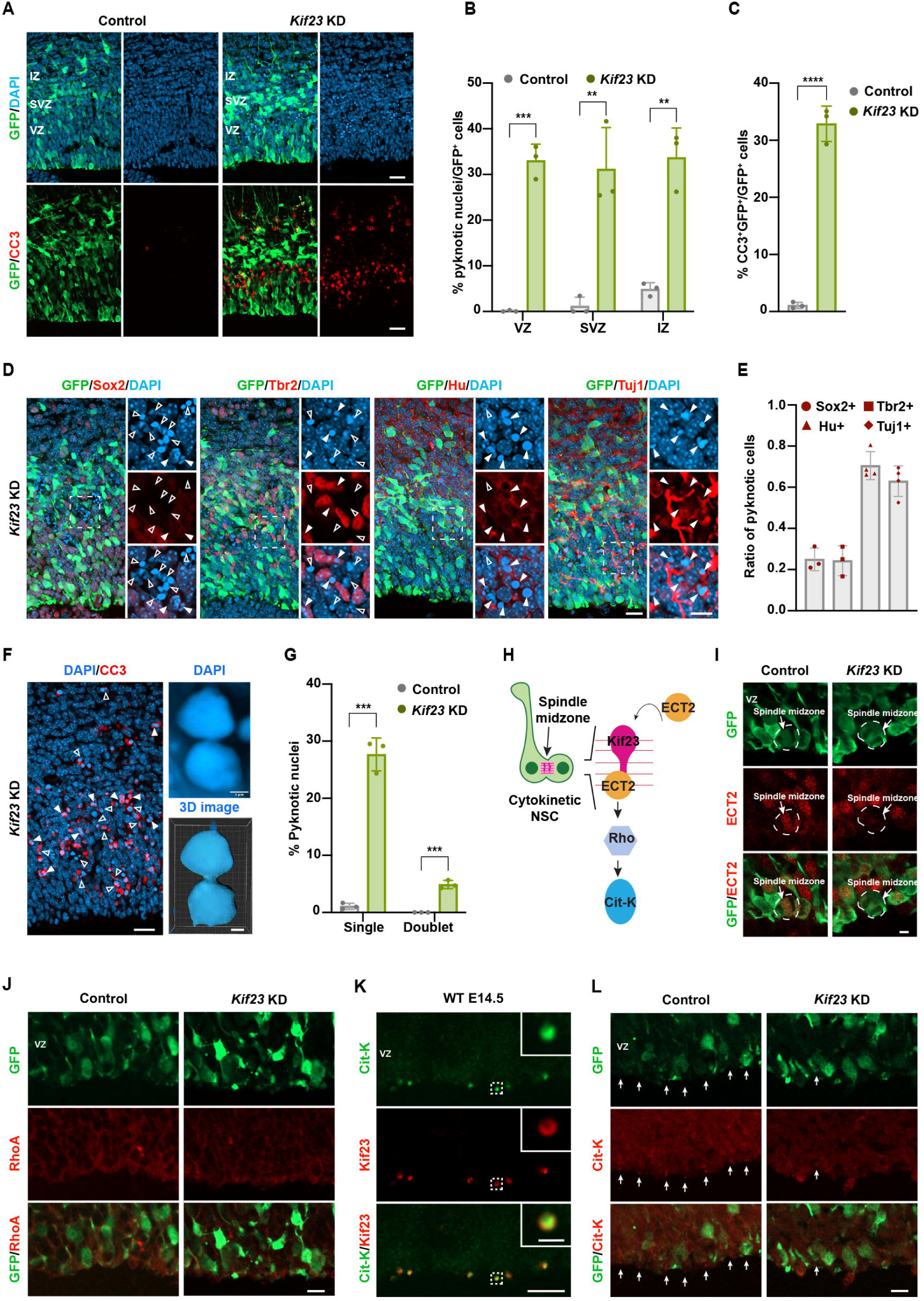
Kif23 deficiency leads to cytokinetic defect and neuronal apoptosis. **(A-C)** Representative images of E15.5 control and *Kif23-*KD cortices stained against GFP, CC3, and DAPI **(A)**. Scale bars, 20 μm. Quantification of the percentage of pyknotic cells **(B)** or CC3^+^ cells **(C)** relative to the total GFP^+^ cells. The data represent the mean ± SD (n = 3). Multiple *t*-tests, \*\**P* < 0.01; \*\*\**P* < 0.001 **(B)**; Two-tailed Student’s *t*-test, \*\*\*\**P* < 0.0001 **(C)**. **(D and E)** Representative images of E15.5 *Kif23-*KD cortical sections stained for GFP, Sox2, Tbr2, Hu, Tuj1, and DAPI **(D)**. Scale bars, 20 μm. Close arrowheads indicate Sox2^+^, or Tbr2^+^, or Hu^+^, or Tuj1^+^ pyknotic cells. Open arrowheads indicate Sox2^−^, or Tbr2^−^, or Hu^−^, or Tuj1^−^ pyknotic cells. Quantification of the ratio of pyknotic cells positive for Sox2, Tbr2, Hu, and Tuj1 **(E)**. The data represent the mean ± SD (n = 3 for Sox2^+^ group; n =3 for Tbr2^+^ group; n = 4 for Hu^+^ group; n =4 for Tuj1^+^ group). **(F and G)** Representative images of E15.5 *Kif23* siRNA transfected cortices stained for CC3 and DAPI **(F)**. Top right panel: binucleated cell stained with DAPI. Bottom right panel: the 3D image of the binucleated cell. Scale bars, 20 μm (left) and 1 μm (right). Open arrowheads indicate pyknotic single with micronuclei. Closed arrowheads indicate pyknotic doublets. Quantification of the percentage of pyknotic cells **(G)**. The data represent the mean ± SD (n = 3). Multiple *t*-tests, \*\*\**P* < 0.001. **(H)** Model of the localization of the cytokinetic molecules from the literature. **(I and J).** Images of E15.5 control and *Kif23-*KD cortical sections stained for GFP and ECT2 **(I)** or GFP and RhoA **(J)**. Examples of cells at the anaphase stage are outlined, and arrows point to the spindle midzone. Scale bar, 5 μm **(I)**; 10 μm **(J)**. **(K)** Images of WT E14.5 cortex to show the co-localization of Cit-K and Kif23. Scale bar, 5 μm. **(L)** Images of E15.5 control and *Kif23-*KD cortical sections stained for GFP and Cit-K. Arrows point to apical midbodies. Scale bar, 10 μm.

To identify the specific cell types undergoing apoptosis, we performed immunostaining with antibodies against various cellular markers. Notably, the pyknotic nuclei were predominantly observed in cells expressing neuronal markers Hu (∼70 %) and Tuj1 (∼60 %). In contrast, only a small proportion of pyknotic nuclei were detected in cells expressing NSPC marker Sox2 (∼25%) and IP marker Tbr2 (∼24%) within *Kif23*-KD cortices (Figures 5D and E). These findings provide a crucial insight, suggesting that differentiating neurons are particularly susceptible to excessive apoptotic cell death following *Kif23*-KD. Further characterization of the apoptotic cells revealed the presence of pyknotic single cells with micronuclei and pyknotic doublets (∼7%) (Figures 5F and 5G). These micronuclei and pyknotic doublets closely resemble binucleated cells, a phenomenon known to result from improper cytokinetic events (Tedeschi et al., 2020; Xie et al., 2021). Consequently, our results put forth the intriguing notion that a subset of cells fails to complete cytokinesis in Kif23-deficient NSPCs, shedding light on the intricacies of Kif23’s role in cell division and survival.

### Molecular mechanism of Kif23-mediated cytokinetic regulation in NSPCs

In our quest to unravel the molecular mechanisms by which Kif23 influences cytokinesis in the developing cortex, we further analyzed the localization of ECT2, a Rho guanine nucleotide exchange factor, in dividing NSPCs (Figure 5H) (Nishimura & Yonemura, 2006). Following *Kif23*-KD, a notable reduction in ECT2 expression within the spindle midzone of dividing NSPCs was observed in comparison to the control group (Figure 5I). Additionally, we explored the localization of RhoA, a key determinant of the division plane (Kamijo et al., 2006), which was found to be disrupted on the apical side of Kif23-depleted cortices (Figure 5J). Furthermore, we uncovered the co-localization of Kif23 with Cit-K, a RhoA effector kinase, within the midbody (Figure 5K). Subsequent analysis revealed a notable reduction in the expression of Cit-K within the apical midbodies under the *Kif23-*KD condition (Figure 5L). These collective findings strongly underscore the pivotal role of Kif23 in orchestrating the proper localization of critical players like ECT2, RhoA, and Cit-K during the intricate processes of cytokinesis in NSPCs.

### Loss of *Kif23* activates cell death and cell cycle arrest pathways

It is well established that binucleated cells are more susceptible to DNA damage when they re-enter the next cell cycle (Hayashi & Karlseder, 2013; S Pedersen et al., 2016). Given our observation of a substantial presence of binucleated cells (Figure 6A), we proceeded to assess DNA damage by staining with the maker γ-H2AX. Notably, within the *Kif23*-KD cortices, we observed a pronounced upregulation of γ-H2AX, which was not evident in the control cortices (Figures 6B and 6C). In the realm of NSPCs, DNA damage is known to activate p53, which, in turn, triggers cell cycle arrest and apoptosis (Shimada et al., 2015; Homma et al., 2021). We further probed whether *Kif23*-KD led to the activation of p53. In comparison to the control group, we noticed a significant increase in the number of p53^+^ cells within *Kif23*-KD cortices (Figures 6D and 6E).

**Figure 6:**
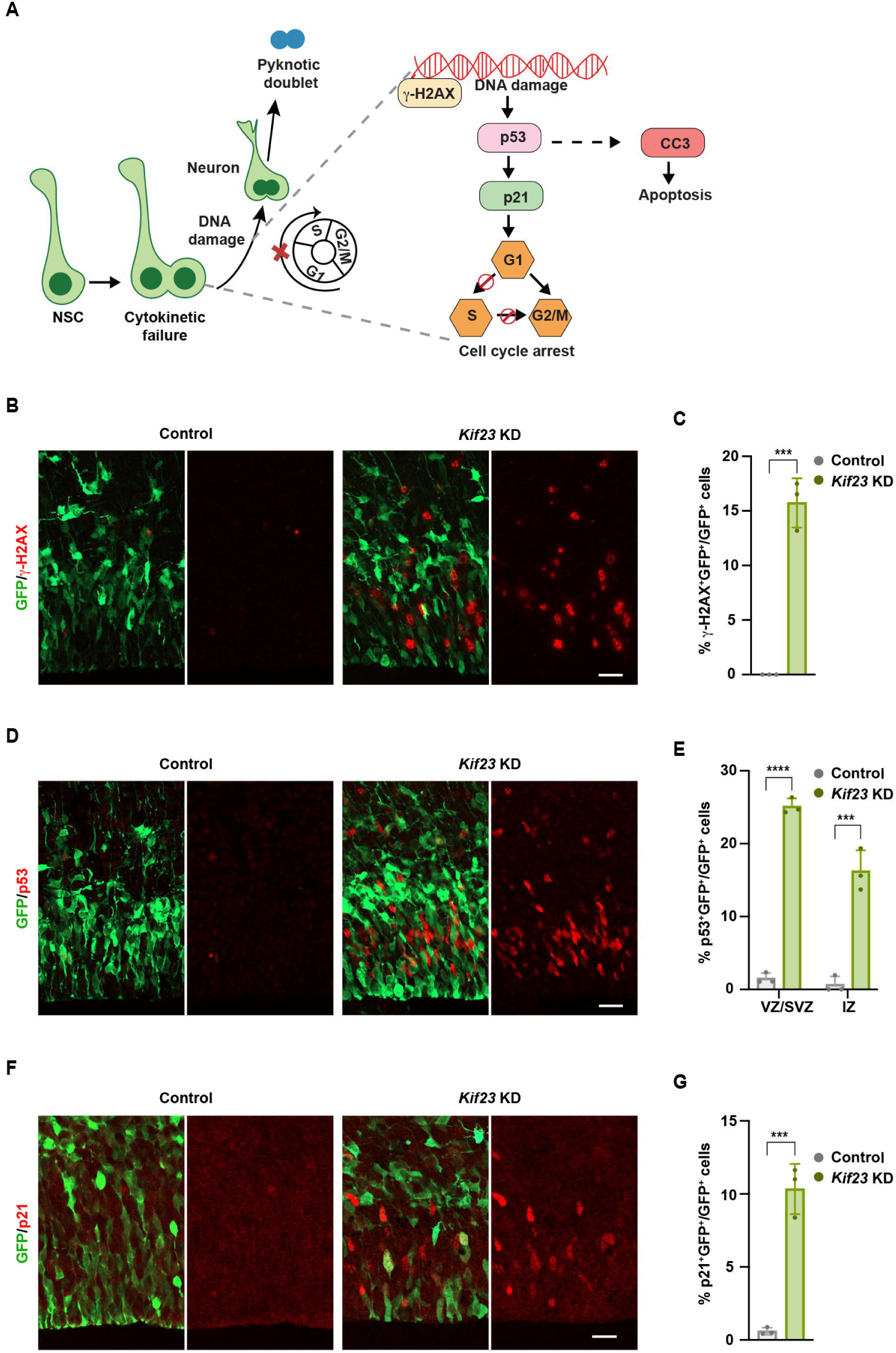
Loss of Kif23 results in upregulation of cell cycle arrest and cell apoptosis pathway. **(A)** Schematic diagram of cell cycle arrest and cell death pathway in the absence of Kif23. **(B-G)** Representative images of E15.5 control and *Kif23-*KD cortices stained for GFP and γH2AX **(B)** or GFP and p53 **(D)** or GFP and p21 **(F)**. Scale bars, 20 μm. Quantification of the percentage of GFP^+^ cells that are γH2AX^+^ **(C)**, p53^+^ **(E)** or p21^+^ **(G)**. The data represent the mean ± SD (n = 3). Two-tailed Student’s *t*-test, \*\*\**P* < 0.001 **(C and G)**; Multiple *t*-tests, \*\*\**P* < 0.001; \*\*\*\**P* < 0.0001 **(E)**.

Given the established role of the cell cycle-dependent kinase inhibitor p21 in promoting progenitor cells to exit the cell cycle and initiate differentiation (Siegenthaler & Miller, 2005), we proceeded to investigate the expression of p21. As anticipated, the results revealed a significant increase in the number of p21^+^ cells within the *Kif23*-KD cortices when compared to the control (Figures 6F and 6G). Consequently, our findings strongly support the notion that Kif23 deficiency not only leads to an increase in apoptosis but also impairs the progression of the cell cycle. This dual impact of Kif23 deficiency on both cell survival and cell cycle progression underscores its multifaceted role in regulating NSPC behavior in the developing cortex.

### Kif23 is essential for maintaining the apical junction structure of NSPCs

Recent research has indicated that microtubules, which constitute the central spindle complex, are implicated in the structural maintenance of adherens junctions in epithelial cells (Meng et al., 2008; Breznau et al., 2015). Moreover, it is known that Kif23 interacts with proteins within the adherens junction (van de Ven et al., 2017). In light of these findings, we embarked on an exploration to determine whether Kif23 plays a vital role in preserving the integrity of the apical junction structure. To do this, we scrutinized the distribution of apical surface molecules, including β-catenin, p120-catenin, and F-actin. Our investigations revealed that *Kif23*-KD cortices exhibited a disruption in the accumulation of these molecules at the apical surface in comparison to the control group (Figure 7A). To gain a comprehensive view of the entire apical surface of NSPCs, we prepared whole mounts of E15.5 *Kif23*-KD cortices and subsequently conducted immunostaining using the apical junctional marker ZO-1 (Figure 7B). Significantly, the size of the apical domain in GFP^+^ cells exhibited a noteworthy increase under the *Kif23*-KD condition (Figures 7C and 7D). Interestingly, this impact was not limited to GFP^+^ cells; the apical domain size of GFP^−^ cells also displayed an increase (Figures 7C and 7D). These findings suggest that Kif23 potentially plays a role in maintaining the integrity of apical junctions in NSPCs and that this role might encompass both cell-autonomous and cell-non-autonomous effects on the apical structure following *Kif23*-KD.

**Figure 7.**
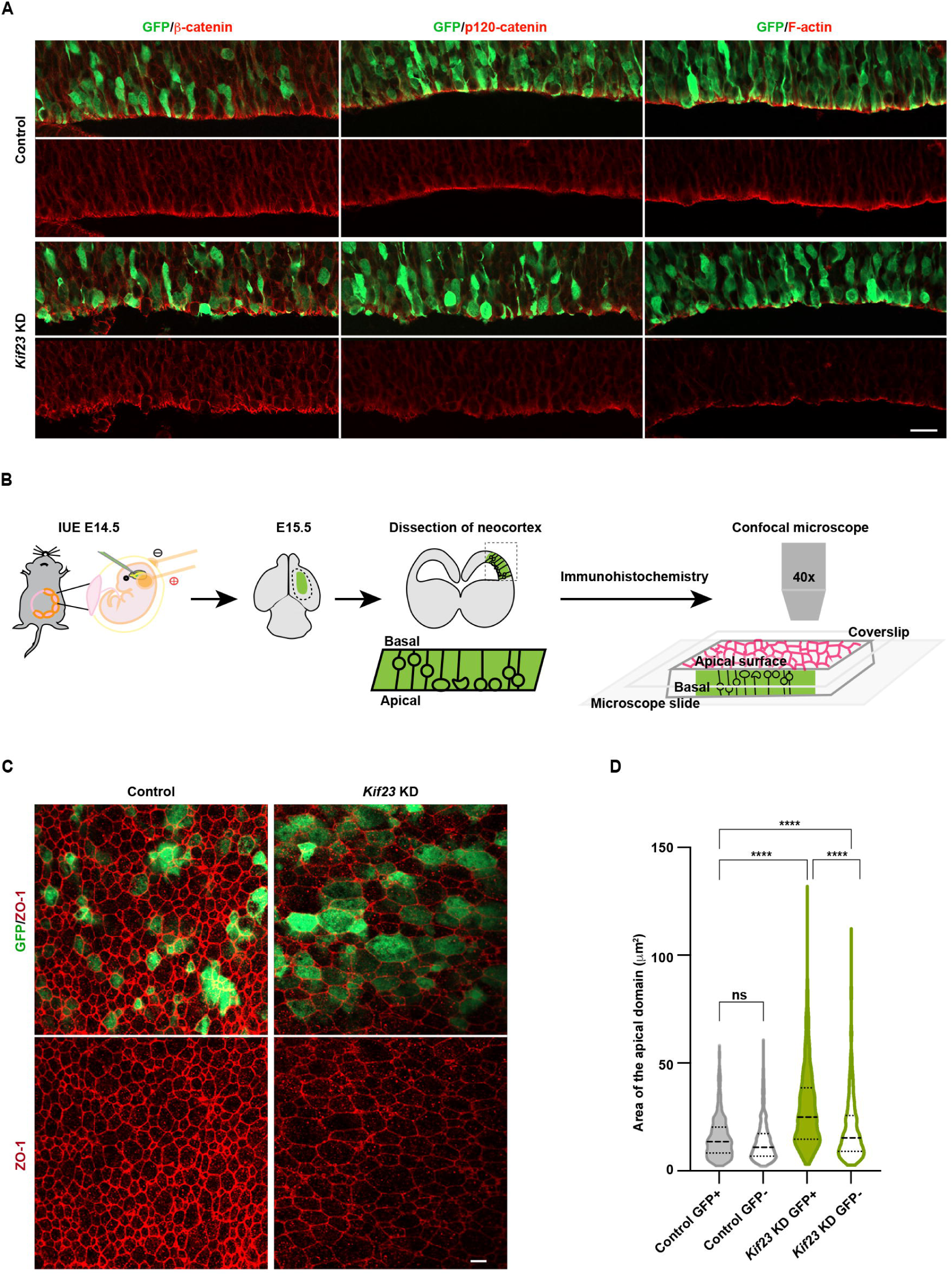
Loss of Kif23 results in the disruption of apical junction integrity. **(A)** Representative images of the control and *Kif23-*KD cortical sections at E15.5 stained for GFP, β-catenin, p120-catenin, and F-actin. Scale bar, 20 μm. **(B-D)** Schematic illustration of the whole mount immunohistochemistry to observe the apical structure **(B)**. Whole-mount images of the electroporated brain sections immunostained for GFP and ZO-1**(C)**. Scale bar, 20 μm. Quantification of the apical domain areas **(D)**. The data represent n = 363 apical domains from 4 samples for the control group; n = 475 apical domains from 6 samples for the *Kif23-*KD group. The thick and thin black horizontal dash lines represent the medians and the quartiles, respectively. One-way ANOVA with Bonferroni’s multiple comparisons test, \*\*\*\**P* < 0.0001; ns: not significant.

### Microcephaly-related *KIF23* mutation fails to rescue *Kif23*-KD phenotypes

To delve deeper into the neuropathology of microcephaly, we initiated our investigation by confirming the expression of *KIF23* using scRNA-seq datasets obtained from human fetal brains during mid-gestation (specifically, gestation week 17-18) (Polioudakis et al., 2019). Our analysis revealed that *KIF23* was notably enriched within the cycling progenitor clusters, specifically those at the S (PgS) and G2/M (PgG2M) phases (Figure 8A). The cells expressing *KIF23* in the human fetal cortex exhibited features similar to those observed in mouse *Kif23* (Figure 1A), underscoring the conservation of Kif23 function in the context of brain development, both in rodents and humans.

**Figure 8.**
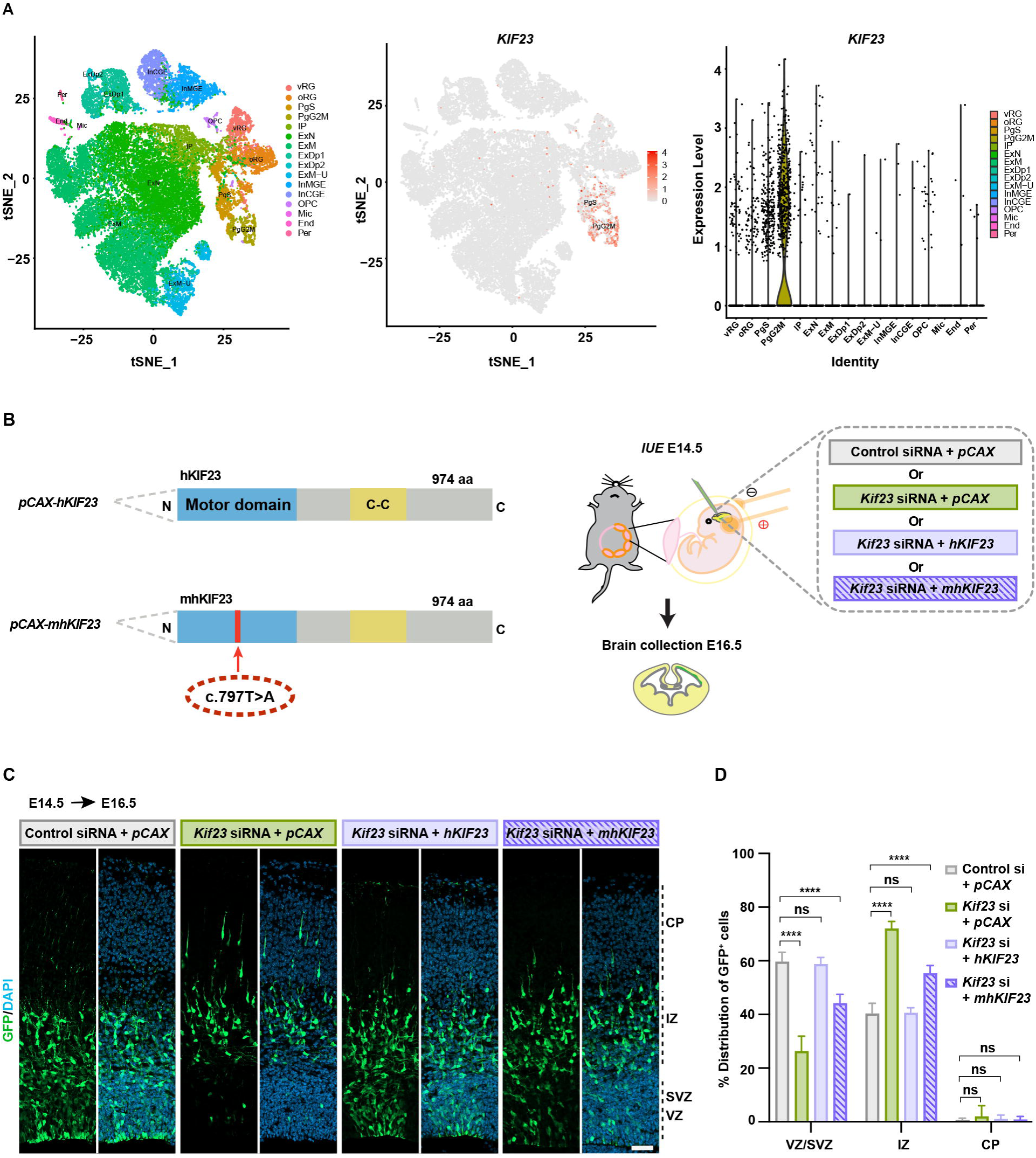
*Kif23* knockdown defects can be rescued by WT human KI23 but not by mutant found in microcephaly patients. **(A)** Single-cell RNA-seq analysis of gestational week 17-18 human fetus cortex showing cell type clusters (left), feature plots (middle), and violin plots (right) to show the expression of *KIF23*. vRG, ventricular radial glia; oRG, outer radial glia; PgS, progenitors in S phase; PgG2M, progenitors in G2M phase; IP, intermediate progenitor; ExN, excitatory newborn neurons; ExM, maturing excitatory neurons; ExM-U, maturing upper layer excitatory neurons; ExDp, deep layer excitatory neurons; InMGE, medial ganglionic eminence interneurons; InCGE, caudal ganglionic eminence interneurons. OPCs, oligodendrocyte precursor cells; End, endothelial cells; Per, pericytes. **(B-D)** Diagram of WT human *KIF23* and mutant *KIF23* constructs, and overview of the experiment **(B)**. Representative images of the mouse cortices two days after *in-utero* electroporation of indicated constructs stained for GFP and DAPI **(C)**. Scale bar, 50 μm. Quantification of the distribution of GFP^+^ cells in VZ/SVZ, IZ, and CP, respectively **(D)**. The data represent the mean ± SD (n = 6 for control siRNA + *pCAX* group; n = 5 for *Kif23* siRNA + *pCAX* group; n = 4 for *Kif23* siRNA + *pCAX-hKIF23* group; n = 8 for *Kif23* siRNA + *pCAX-mhKIF23* group). One-way ANOVA with Bonferroni’s multiple comparisons test, \*\*\*\**P* < 0.0001; ns: not significant.

Next, we asked about potential means of a mutation related to microcephaly by rescuing *Kif23*-deficient condition. We employed human *KIF23* (*hKIF23*) and a variant carrying a missense mutation within the motor domain of *KIF23* (c.755T>A; p.L266H) (referred to as *mhKIF23*). This specific mutation has been identified in patients with microcephaly in a homozygous condition (Karaca et al., 2015). To assess the rescue potential, we co-electroporated *hKIF23* along with *Kif23*-siRNA into mouse embryos at E14.5 and collected the brains at E15.5 to perform immunostaining. The outcome revealed that *Kif23* siRNA effectively reduced the amount of mouse Kif23 protein, while human KIF23 protein remained unaffected (Figures S2A and S2B). This was attributed to the resistance of *hKIF23* cDNA to *Kif23*-siRNA.

Subsequently, we investigated the distributions of GFP^+^ cells at E16.5 in brains co-electroporated with *hKIF23* and *Kif23*-siRNA at E14.5. Under this condition, the distribution of GFP^+^ cells closely resembled that of the control group, indicating the successful rescue of *Kif23*-KD by *hKIF23* (Figures 8B-8D). In contrast, the co-electroporation of *mhKIF23* and *Kif23*-siRNA resulted in a decrease in the proportion of GFP^+^ cells within the VZ/SVZ and an increase in the IZ, as compared to the control and *hKIF23* groups (Figures 8B-8D). These results provide a strong indication that *mhKIF23* was unable to rescue the defects in cortical development induced by *Kif2*3-KD. This observation strongly implies that mutations within the *KIF23* gene can result in a loss of function, which, in turn, can lead to the onset of microcephaly, thus furthering our understanding of the genetic basis of this neurodevelopmental disorder.

## Discussion

In this study, we delved into the dynamic localization and developmental transition of the motor protein Kif23 in NSPCs and its critical roles in multiple processes of cytokinesis during early corticogenesis in mice. Our research uncovered that Kif23 exhibited dynamic localization patterns that change as NSPCs progress through different phases of the cell cycle. Notably, it was consistently present in the spindle microtubule throughout the mitotic stages of NSPCs.

As a general principle, the orientation of the spindle cleavage planes has far-reaching consequences in determining the mode of cell division. Spindle cleavage planes oriented vertically in relation to the apical surface tend to yield symmetric division, generating two proliferative progenitors, while horizontally or obliquely oriented spindle cleavage planes result in asymmetric division, producing one proliferative daughter cell and one postmitotic neuron (Haydar et al., 2003; Noctor et al., 2004). Our findings demonstrated that the disruption of Kif23 function in the developing cortex led to abnormalities in spindle organization and orientation, manifesting as an increased prevalence of NSPCs with asymmetric spindle cleavage planes. These observations are in line with prior studies implicating mutations in several centrosomal genes like *Nde1*, *Cdkrap2*, and *PPP4c*, which similarly induce spindle orientation defects. This, in turn, results in an increase in asymmetric mitotic cleavage and the premature exit of NSPCs from the cell cycle (Feng & Walsh, 2004; Xie et al., 2013). Therefore, our study postulated that the precocious neurogenesis observed by *Kif23*-KD, is likely mediated by spindle orientation defects.

Improper cytokinesis can manifest in two distinct outcomes: 1) binucleation, which arises from cleavage furrow regression, and 2) the formation of individual cells with micronuclei, a consequence of premature chromatin bridge resolution (Gromley et al., 2003; Pampalona et al., 2012). Our research underscores that deficiencies in Kif23 function precipitated both of the phenotypes: the presence of doublet pyknotic cells and single pyknotic cells with micronuclei, ultimately culminating in reduced neuronal production. Consequently, our findings suggest the indispensable role of Kif23, which extends beyond its involvement in regulating the mitotic spindle to encompass the pivotal task of ensuring proper cytokinesis.

Our investigation unveiled that *Kif23*-KD had a notable impact on the expression of cytokinetic molecules within NSPCs, including ECT2, RhoA, and Cit-k. Recent research in *Drosophila* has shed light on the necessity of Kif23 for the transportation of ECT2 as cargo to the spindle midzone and the equatorial cell cortex (Warecki & Tao, 2023). However, to comprehensively understand the underlying mechanisms, further study is warranted to ascertain whether Kif23 physically interacts with ECT2, RhoA, and Cit-k within NSPCs. Additionally, investigating whether these molecules collaborate in tandem or operate independently in the regulation of cytokinesis represents an intriguing avenue for future exploration.

The molecular mechanisms underlying the detection of cytokinetic abnormalities and initiation of cell apoptosis in the developing brain are not well understood. Previous studies have indicated that when cells are arrested in the G1 phase, they tend to exit the cell cycle rather than undergo apoptosis, both in cultured cells and in the fly brain (Ganem et al., 2014; Gogendeau et al., 2015). In our study, we identified an augmented DNA damage response and increased p53 signaling in the *Kif23*-KD cortices. Additionally, we observed p53 accumulation in both NSPCs and neurons, alongside elevated p21 levels after *Kif23*-KD. This elevated p53 and p21 may account for the premature neuronal differentiation observed in *Kif23-*KD cortices. Since p53 was detected in both NSPCs and neurons following *Kif23*-KD, it suggests that p53 accumulation initiates in NSPCs due to cytokinesis failure in response to DNA damage and persists in the differentiated neurons, ultimately triggering neuronal apoptosis. Therefore, our findings propose a mechanism linking cytokinesis failure, premature cell cycle exit, and subsequent apoptosis induced by Kif23 dysfunction.

Kif23 is found in the midbodies located on the apical surface of the embryonic mouse cortex (Maliga et al., 2013). Symmetrically dividing NSPCs release their midbodies into the ventricular fluid after cytokinesis (Dubreuil et al., 2007), as observed in our samples stained with Kif23 (Figure 1E). Kif23 expression was notably higher during early neurogenesis, a phase when NSPCs predominantly undergo symmetric division, suggesting a possible association between Kif23-containing midbody release and symmetric division (Figure 1E). In HeLa cells, Kif23 facilitates the localization of midbody-enriched mRNA cargo, leading to translational availability after the M/G1 transition, with implications for cell fate determination, cell division, and proliferation (Park et al., 2023). This raises the intriguing possibility that Kif23, functioning as a motor protein, might transport cell fate determinants within the NSPCs, influencing the fate of their daughter cells.

We observed a significant deterioration in the structure of apical endfeet in *Kif23*-KD cortices, highlighting another crucial role of Kif23 in supporting NSPCs. A study in *C. elegans* revealed that the dynamic localization of the adhesion complex during cytokinesis plays a pivotal role in preserving proper cell contacts and maintaining the integrity of apical surface architectures (Bai et al., 2020). It is conceivable that the loss of Kif23 may lead to a reduction in the expression of junctional proteins such as β-catenin, p120-catenin, and F-actin, both within the NSPCs and in neighboring cells. Importantly, disruptions of the apical structure of NSPCs are well-documented to trigger premature neuronal differentiation (Tamai et al., 2007; Shinohara et al., 2013; Yoon et al., 2017).

Microcephaly can arise from reduced NSPC proliferation, increased premature neuronal differentiation, and elevated cell death (Lizarraga et al., 2010; Chen et al., 2014). Using a public database of human fetal cerebral cortex, we identified robust expression of *KIF23* in dividing NSPCs (Polioudakis et al., 2019), suggesting its potential involvement in human brain development. In the present study, we assessed the functionality of human KIF23 through a *Kif23-*KD model in the mouse cortex. Our results demonstrated that WT *hKIF23* effectively rescued the phenotypes of *Kif23-*KD, whereas a motor domain mutant (*mhKIF23*) did not. This strongly implies that mutations in *KIF23* gene are likely responsible for the neuropathology associated with human microcephaly.

In summary, our findings underscore the critical importance of proper motor protein function in cytokinesis of NSPCs and the preservation of their apical structure in early development of the mammalian cortex. Further investigations encompassing human genetics and animal models promise to unveil the intricate molecular and cellular mechanisms underpinning the fascinating process of brain development.

## Methods

### Data processing of Visium spatial gene expression profiles

Spatial transcriptome analysis (Visium, 10x Genomics) was performed on the brain sample of WT E15.5 mice as described previously (Tsai et al., under revision). Briefly, sagittal brain sections (10-μm thick) were placed on gene expression (GE) slides (10X Genomics). The captured cDNA library tagged with spatial barcode were sequenced, and the resulting data was analyzed by Space Ranger software (Version 1.2.1). Gene alignment was performed using the mouse reference genome (MM10), and Loupe Browser 6.4.0 was used for data analysis. *Kif23* expression was determined based on unique molecular identifier (UMI) counts in each spot, and violin plots were used to display *Kif23* expression in each cluster.

### Data processing of scRNA-seq analysis

All scRNA-seq data used in this study are available from the public databases: mouse cortex: GSE123335 (Loo et al., 2018), and human cortex: phs001836 (Polioudakis et al., 2019; data was obtained from http://geschwindlab.dgsom.ucla.edu/pages/codexviewer). Normalization, dimensionality reduction, clustering, and visualization of scRNA-seq data were performed by Seurat (v4.3, Satija et al., 2015) function of NormalizeData, RunPCA, RunUMAP, FindNeighbors, FindCluster with default parameters.

### Animals

C57BL/6J mice (CLEA Japan) were used as wild-type (WT) in this study. Embryonic day 0.5 (E0.5) was defined as midday on the day when a vaginal plug was detected. All experiments were carried out in compliance with the National Institutes of Health guidelines for the care and use of laboratory animals, and experimental procedures were approved by the Ethics Committee for Animal Experiments of Tohoku University Graduate School of Medicine (2019 MdA-018-07).

### *In situ* hybridization

The dorsal telencephalon of WT E14.5 mice was collected and fixed with 4 % paraformaldehyde for 16 hours. *In situ* hybridization on the frozen section was performed as previously described (Kikkawa et al., 2013). cDNA fragment encoding *Kif23* (NM_024245, nucleotides 866-1367) was cloned into pBluescript II SK (–) (Stratagene). Digoxigenin-labeled RNA probe was synthesized using DIG RNA labeling kit (Roche). BZ-9000 fluorescence microscopy system (KEYENCE) was used to capture images.

### Immunohistochemistry

Immunohistochemistry was performed as described previously (Kikkawa et al., 2013). The frozen brain sections were incubated with primary antibodies diluted with 3% bovine serum albumin (BSA) in Tris-buffered saline with 0.1% Triton X-100. The primary antibodies were used as follow: rabbit anti-Kif23 (1:1000; Invitrogen), chicken anti-GFP (1:1000; Abcam), rabbit anti-Pax6 (1:1000; MBL), rabbit anti-Tbr2 (1:1000; Abcam), rabbit anti-Ki67 (1:500; Abcam), rabbit anti-p53 (1:1000; Leica Biosystems), rabbit-anti-active caspase-3 (1:1000; BD bioscience), rat anti-Tbr2 (1:1000; Invitrogen), mouse anti-β catenin (1/1000; BD bioscience), mouse anti-p120 catenin (1/1000; BD bioscience), mouse anti-p21 (1:1000; Proteintech Group), mouse anti-phospho-histone H3 (1/1000; Cell signaling), mouse anti-Tuj1(1/1000; BioLegend), mouse anti-HuC/D (1:200; Molecular Probe), mouse-anti α-tubulin (1:200, Sigma), mouse-anti γ-tubulin (1:300, Sigma), and mouse-anti γ-H2AX (1:1000; Millipore). Secondary antibodies were as follows: Cy3-conjugated affinity purified anti-rabbit or anti-mouse IgG donkey antibodies (1: 500; Jackson Immunoresearch Laboratories), and Alexa 488-conjugated affinity-purified anti-mouse IgG or anti-chicken IgY goat antibodies (1: 500; Invitrogen). 4′,6-Diamidino-2-phenylindole dihydrochloride (DAPI) (1:1000; Sigma) was used as nuclear counterstaining. All images were acquired using a confocal laser microscope Zeiss LSM800 (Carl Zeiss).

### *In-utero* electroporation

*In-utero* electroporation (IUE) was performed as described previously with slight modification (Nagai et al., 2022). Pregnant WT mice (E14.5) were anesthetized with isoflurane. A mixture of siRNA and *pCAG-EGFP* (final concentration of 2 μg/μl and 0.5 μg/μl, respectively) was injected into the lateral ventricle of the brain by using a glass capillary with the electric microinjector system (IM-300, Narishige). Five electric pulses at 40 V with a duration of 50 ms per pulse at 1s intervals were applied through the uterus using forceps-type electrodes (LF650P5, BEX) connected to an electroporator (CUY-21, BEX).

### Construction of human *KIF23* plasmid

Human *KIF23* cDNA was synthesized based on NCBI sequencing data (accession no. NM_001367805) (eurofins). The ORF of human *KIF23* was cloned within *pCAX* vector using In-Fusion system (TaKaRa). The disease associated c.755T>A mutation was introduced to *human KIF23* cDNA by the PCR-based mutagenesis as described (Xia et al., 2015). Instead of Phusion DNA polymerase, KOD plus DNA polymerase (TOYOBO, Tokyo, Japan) was used. The mutation was confirmed by the Sanger sequencing.

### EdU labeling

EdU labeling was performed as described previously with slight modification (Ji et al., 2017). 5-Ethynyl-2’-deoxyuridine (EdU) (50 mg/kg) was injected intraperitoneally into pregnant mice at the gestational day 15 or 16. For cell proliferation analysis, regnant mice were sacrificed 2 hours after the injection at the gestational day 16. For cell cycle exit analysis, the pregnant mice were sacrificed 24 hours after the injection. EdU immunostaining was performed according to the Click-iT^TM^ Plus EdU Alexa Fluor^TM^ 555 imaging kit protocol (Invitrogen).

### Whole mount immunohistochemistry

Whole mount immunohistochemistry was performed as described previously with minor modifications (Shinohara et al., 2013). Electroporated embryos at E15.5 were perfused with 4% paraformaldehyde, and their brains were isolated and fixed in the same fixative for 25 mins. Dorsolateral parts of the cerebral cortex were dissected, fixed for 2 mins, and incubated in a blocking solution (3% BSA) for 1 hour at room temperature. The samples were then incubated overnight at 4°C with primary antibodies, including chicken anti-GFP (1:1000; Abcam) and rabbit anti-ZO-1 (1/1000; Invitrogen). After incubation with secondary antibodies, whole mounts were cover-slipped with VECTASHIELD® antifade mounting medium (Vector Laboratories).

### Spindle orientation measurement

Spindle orientation measurement was done following the protocol described previously (Yingling et al., 2008). Briefly, the angle θ of the mitotic cleavage plane of cells in late metaphase or anaphase was measured relative to the surface plane of the ventricular zone. The angle α of spindle orientation was calculated as 90 degrees minus the angle θ. Analysis was performed by using the measurement tools of the Fiji software (National Institute of Health).

### Statistical analysis

The cell counts were carried out using Fiji software (National Institute of Health). Statistical analyses were conducted using GraphPad Prism (version 9.5.0), including Student’s *t*-tests, multiple unpaired *t*-tests, one-way ANOVA with Bonferroni’s multiple comparisons test, and two-way ANOVA with Bonferroni’s multiple comparisons test. Differences with p<0.05 were considered statistically significant. *P* values are denoted as follows: \**p*<0.05, \*\**p*<0.01, \*\*\**p*<0.001, \*\*\*\**p*<0.0001.

## Acknowledgments

We would like to thank Nobutaka Hirokawa, Yoshio Wakamatsu, Hiroshi Shinohara, and Shohei Ochi for their technical advice and valuable comments. We are grateful to Ms. Sayaka Makino for animal care and technical support. We thank all other members of the Osumi laboratory for their valuable comments and discussion. This work was supported by JSPS KAKENHI funding (#18K14999, 23K06297) to T.K and AMED (#JP21wm0425003) to N.O.

## Author Contributions

Conceptualization: S.N, T.K, and N.O; Methodology & Investigation: S.N, T.K, K.I, S.M, S.N and J.W.T; Writing-Original Draft: S.N, T.K, J.W.T, and N.O; Writing-Review & Editing: S.N, T.K, K.I, S.M, M.H; K.T; S.N, J.W.T, and N.O; Supervision: T.K, J.W.T, and N.O; Funding acquisition: T.K, and N.O.

## Declaration of Interests

The authors declare no competing interests.

## Notes

### Competing Interest Statement

The authors have declared no competing interest.

