## Supplementary information for "Kinesin family member Kif23 regulates cytokinetic division and maintains neural stem/progenitor cell pool in the developing neocortex"

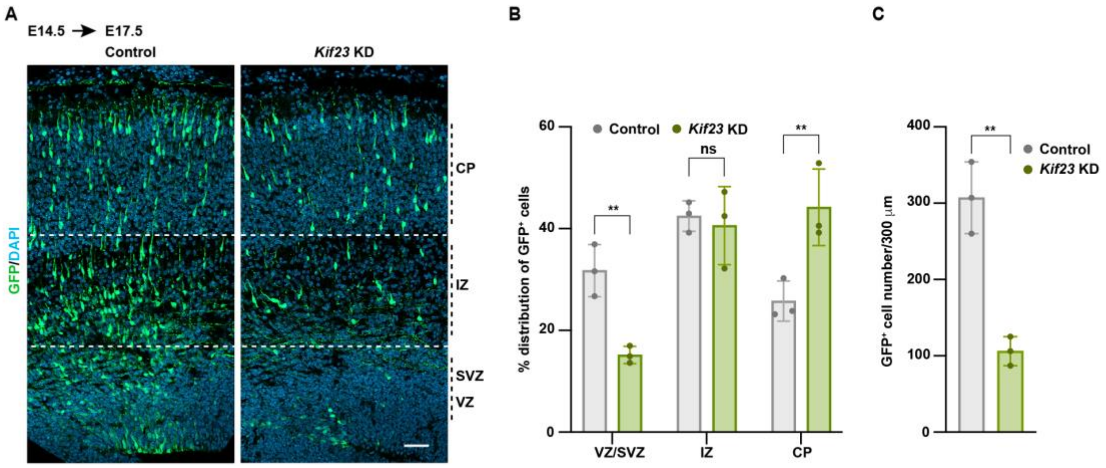


**Supplementary Figure 1: Loss of Kif23 leads to profound loss of GFP expressing cells**

**(A)** Representative images of $GFP^+$ cells distribution in the control and *Kif23*-KD cortices at E17.5. Dashed lines illustrate the borders among VZ/SVZ, IZ, and CP. Scale bar, 50 μm.

**(B)** Quantification of $GFP^+$ cells distribution in VZ/SVZ, IZ, and CP, respectively. The data represent the mean ± SD (n = 3). Two-way ANOVA with Bonferroni's multiple comparison test, **P < 0.01; ns: not significant.

**(C)** Quantification of the average number of $GFP^+$ cells. The data represent the mean ± SD (n = 3). Student's t-test, **P < 0.01; ns: not significant.

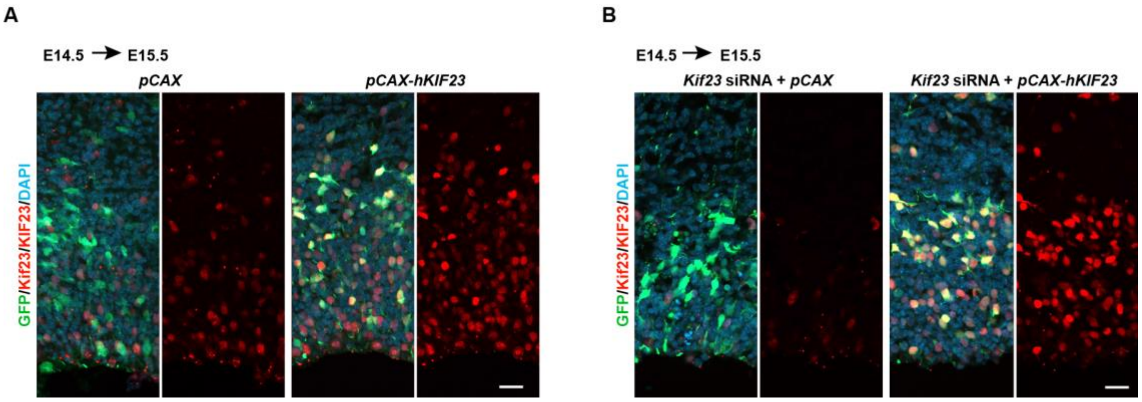


**Supplementary Figure 2: *Kif23* siRNA decreases mouse Kif23 protein levels, not human KIF23**

**(A)** Representative images of mouse cortices one day after electroporation of *pCAX* or *pCAX-hKIF23* stained for GFP and Kif23/KIF23. Scale bar, 20 μm.

**(B)** Representative images of mouse cortices one day after electroporation of *Kif23* siRNA/ *pCAX* or *Kif23* siRNA/*pCAX-hKIF23* stained for GFP and Kif23/KIF23. Scale bar, 20 μm.
